# Predicting causal variants affecting expression using whole genome sequence and RNA-seq from multiple human tissues

**DOI:** 10.1101/088872

**Authors:** Andrew Anand Brown, Ana Viñuela, Olivier Delaneau, Tim Spector, Kerrin Small, Emmanouil T Dermitzakis

## Abstract

Genetic association mapping produces statistical links between phenotypes and genomic regions, but identifying the causal variants themselves remains difficult. Complete knowledge of all genetic variants, as provided by whole genome sequence (WGS), will help, but is currently financially prohibitive for well powered GWAS studies. To explore the advantages of WGS in a well powered setting, we performed eQTL mapping using WGS and RNA-seq, and showed that the lead eQTL variants called using WGS are more likely to be causal. We derived properties of the causal variant from simulation studies, and used these to propose a method for implicating likely causal SNPs. This method predicts that 25% - 70% of the causal variants lie in open chromatin regions, depending on tissue and experiment. Finally, we identify a set of high confidence causal variants and show that they are more enriched in GWAS associations than other eQTL. Of these, we find 65 associations with GWAS traits and show examples where the gene implicated by expression has been functionally validated as relevant for complex traits.

---

Genetic association mapping produces statistical links between phenotypes and genomic regions, but identifying the causal variants themselves remains difficult. Complete knowledge of all genetic variants, as provided by whole genome sequence (WGS), will help, but is currently financially prohibitive for well powered GWAS studies. To explore the advantages of WGS in a well powered setting, we performed eQTL mapping using WGS and RNA-seq, and showed that the lead eQTL variants called using WGS are more likely to be causal. We derived properties of the causal variant from simulation studies, and used these to propose a method for implicating likely causal SNPs. This method predicts that 25% − 70% of the causal variants lie in open chromatin regions, depending on tissue and experiment. Finally, we identify a set of high confidence causal variants and show that they are more enriched in GWAS associations than other eQTL. Of these, we find 65 associations with GWAS traits and show examples where the gene implicated by expression has been functionally validated as relevant for complex traits.

Genome-wide associations studies (GWAS) have uncovered 1,000s of genetic associations between regions of the genome and complex traits (Welter *et al.*, 2014), but moving from the association to identifying the mechanism behind it has proven complicated (Spain and Barrett, 2015). A first step would be to identify the exact variant behind the association, as exact localisation would allow exploration as to which transcription factor binding sites and regulatory elements are affected. This, however, is complicated by the fact that most loci tested in GWAS studies are not directly measured, but instead imperfectly imputed (Marchini and Howie, 2010). Whole-genome sequence (WGS) data does directly ascertain all genotype calls, but despite falling costs it is still very expensive on the sample sizes of modern GWAS studies (Supplementary Table S1). In contrast, studies looking at genetic variants and gene expression (eQTL studies) have discovered 1,000s of associations using few hundreds of samples, a scale at which collecting whole genome sequence data is feasible (Lappalainen *et al.*, 2013).

In this work we describe analysis combining for the first time two previously published datasets derived from individuals in the TwinsUK cohort: RNA-seq from four tissues (Brown *et al.*, 2014; Buil *et al.*, 2015) and WGS from the UK10K project (UK10K Consortium *et al.*, 2015). We explore the properties of causal variants using simulations, leading us to propose the CaVEMaN method (Causal Variant Evidence Mapping using Non-parametric resampling), which estimates the probability that a particular variant is causal. Application of this method allows us to produce a robust set of likely causal SNPs; this could be an important resource for developing methods to call personalised regulatory variants from whole-genome sequence and sequence annotations.

In whole genome sequence every variant is directly measured, the degree to which this increases power to map eQTLs by removing noise from imputation errors is currently unknown. For a simple comparison, we mapped independent eQTLs within 1Mb of the transcription start site for protein coding genes and lincRNAs in four tissues (fat, lymphoblastoid cell lines (LCLs), skin and whole blood) using individuals for which expression, sequence and genotype array data were all available (N from 242 (whole blood) to 506 (LCLs)). Using an eQTL mapping strategy based on stepwise linear regression, we identify 27,659 independent autosomal eQTLs affecting 11,865 genes using whole genome sequence (8,690,715 variants), and 26,351 affecting 11,642 genes using genotypes called from arrays and imputed into the 1000 Genomes Project Phase 1 reference panel (6,263,243 variants) (Figure 1, an analysis of all individuals with expression and WGS data (N from 246-523) and including the X chromosome found 28,141 eQTLs affecting 12,243 genes). This means just a 3.7% increase in discovered eQTLs using WGS; balanced against at least a ten-fold increase in cost of collecting the data, it does not seem a worthwhile exercise yet.

**Figure 1:**
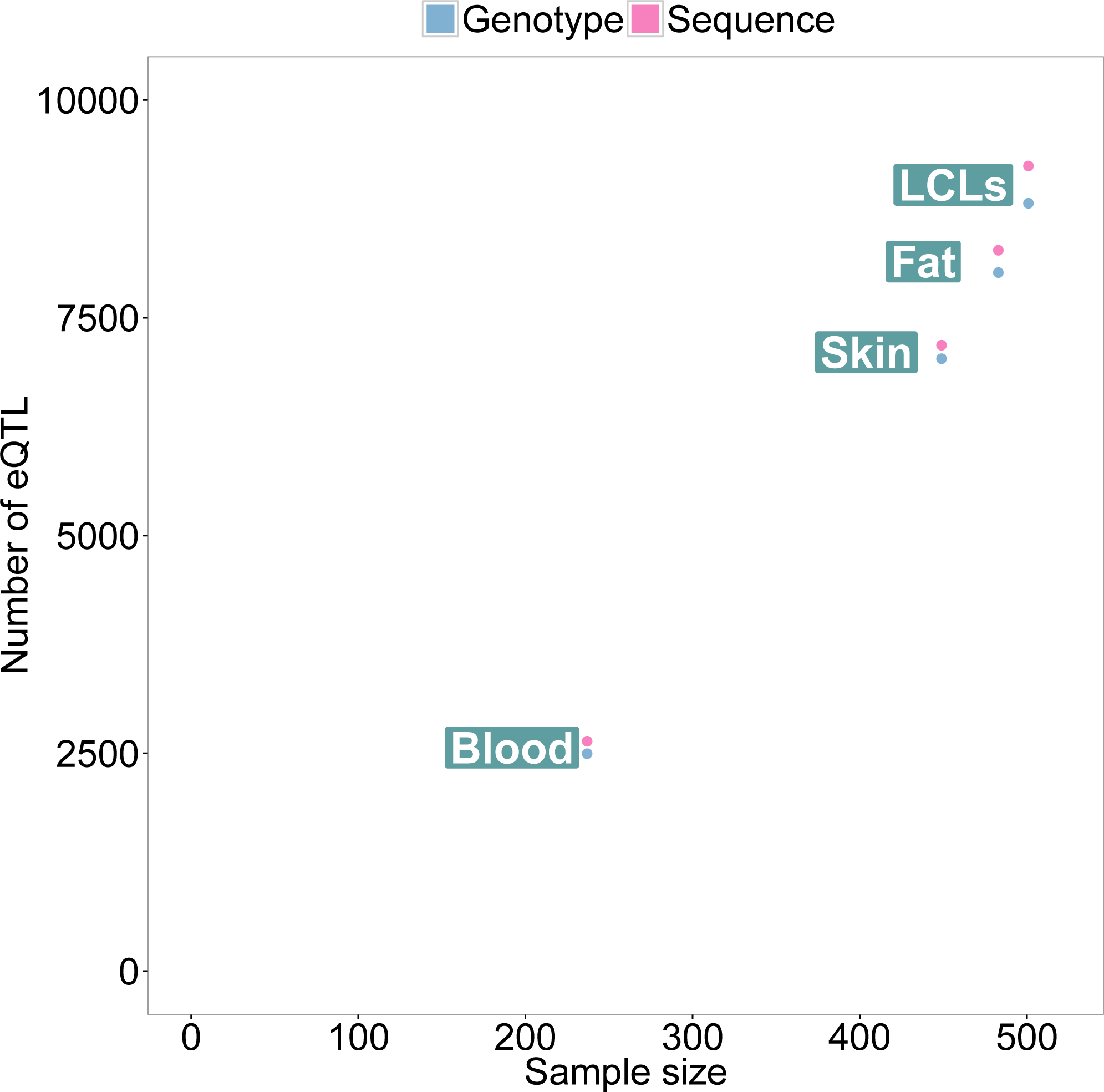
Number of autosomal eQTLs discovered in each tissue when genotype information is provided by arrays imputed into a reference panel and by whole genome sequencing. There is a modest (3.7%) increase in the number of eQTL discovered with WGS.

We frequently observe that the lead eQTL variant (LEV, by which we refer to the variant most associated with the trait) differs between the two datasets. As genotypic uncertainty should be reduced for WGS, due to lack of imputation biases, we expect the WGS LEVs to be the causal variant more frequently than LEVs from genotype arrays. To test this hypothesis, we looked for enrichment of WGS-derived LEVs relative to array-genotype-derived in biochemically active regions of the genome. Indeed, for 30 out of 31 experiments carried out by the Roadmap Epigenomics consortium (Roadmap Epigenomics Consortium *et al.*, 2015) in relevant tissues, we see significant enrichment of sequence LEVs compared to genotype LEVs falling in DNase1 hypersensitivity sites (DHS) (Odds ratio, 1.17-1.40, Figure 2). From this we infer that the LEVs called with sequence are more likely the causal variant.

**Figure 2:**
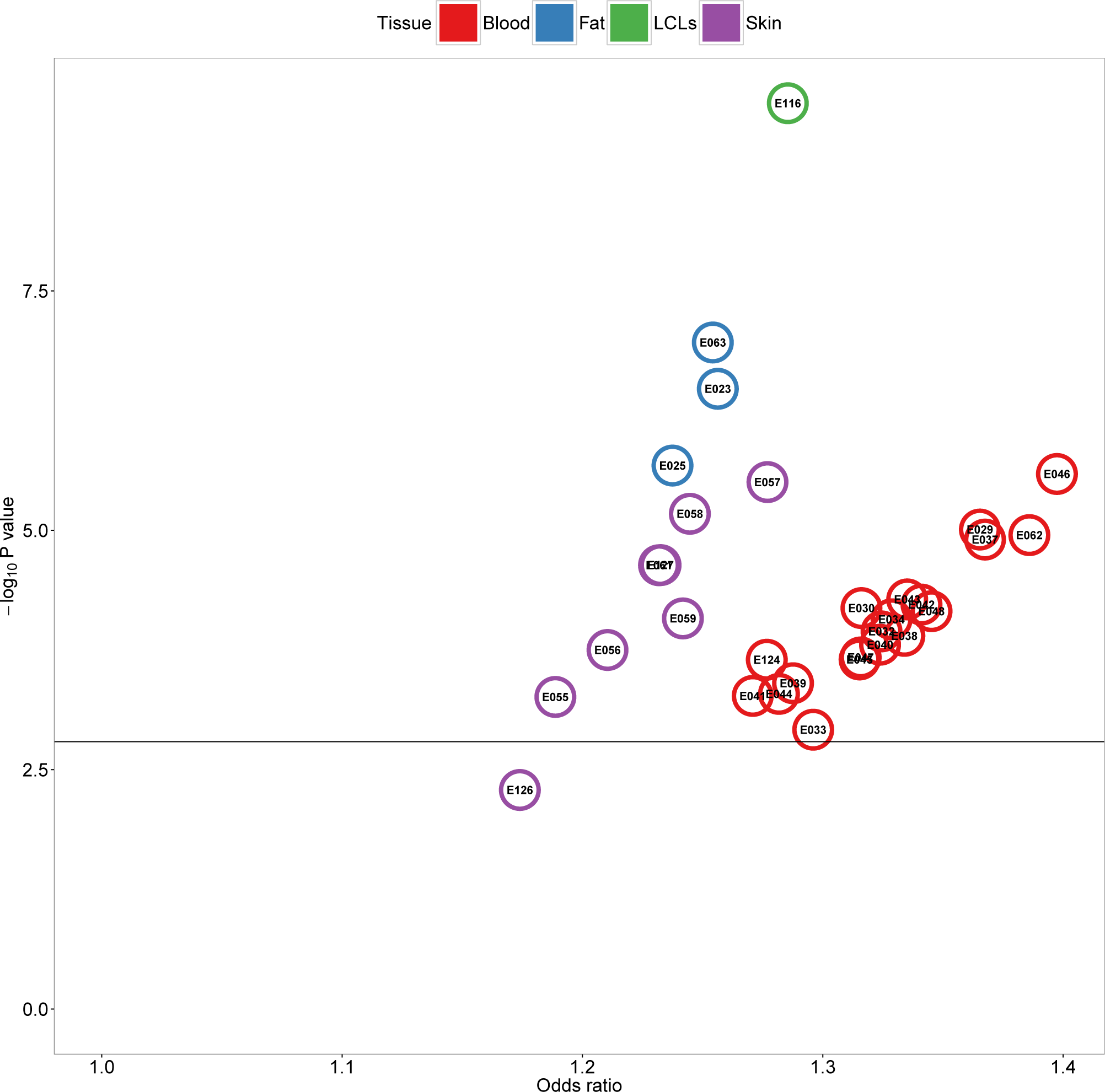
Odds ratio and P value for enrichment of lead eQTL variant called from sequence being located in DNase hypersensitivity sites (Roadmap Epigenomics Consortium *et al.*, 2015) relative to LEVs called from array derived genotypes. A total of 31 experiments related to the tissue from which RNA-seq was collected were analysed, the code given relates to the Roadmap Epigenomics code, Supplementary Table S2 lists the original experiment. All but enrichment of skin eQTL in DHS assyed in NHDF-Ad Adult Dermal Fibroblast Primary Cells were Bonferroni significant (P < 0.05).

To better understand properties of causal variants we simulated expression datasets where the causal variant is known, with properties matched to those of the LEVs from the original eQTL mapping with sequence genotypes (considering effect size, distance to the transcription start site and minor allele frequency). Repeating the eQTL mapping on these simulated datasets, we found that in 45% of cases the causal variant was the LEV. This number was consistent across tissues, despite sample size and power to map eQTLs being much reduced for whole blood (Supplementary Figure S1). This number is also similar to that obtained from the analysis of the Geuvadis data (55%), which used a different methodology based on difference in P values and enrichment in DHS. We also see a rapid decline when looking at lower ranked candidate variants, with the 10th most associated SNP being only causal in 1% of cases.

Our simulations show that across all genes, the LEV is a strong candidate for the causal variant. However, when considering specific LEVs, causality for that variant will depend on the linkage disequilibrium structure around the true causal variant and phenotypic uncertainty for the expression of the gene of interest. For these reasons we developed the CaVEMaN method, which uses bootstrap methods (Visscher *et al.*, 1996; Lebreton and Visscher, 1998) to estimate the probability that the LEV is the causal variant (see Supplementary Methods for methodological details).

We have applied the CaVEMaN method to all four tissues and the Geuvadis LCL RNA-seq data (N = 445, results in Supplementary File 1). The distributions of probabilities that LEVs are causal are similar across tissues and studies (Figure 3). For 7.5% of the eQTLs the LEV has P > 0.8 of being the causal variant, we refer to these as High Confidence Causal Variants (HCCVs). For comparison, we applied the Caviar method (Hormozdiari *et al.*, 2014) to the largest dataset (TwinsUK LCLs), restricting the analysis to all genes with only one eQTL to remove differences related to inferring presence of multiple eQTLs. Caviar, along with with equivalent Bayesian methods (Chen *et al.*, 2015; Benner *et al.*, 2016; Servin and Stephens, 2007), have previously been suggested as fine-mapping methods for estimating credible sets of SNPs with a given probability of containing the causal variant. There was good agreement on the causal probabilities of the LEV (spearman *ρ* = 0.856, *P* < 10^−216^, Supplementary Figure S3), but the Caviar method produced more conservative estimates of the causal probabilities (median probability 0.12 vs 0.29).

**Figure 3:**
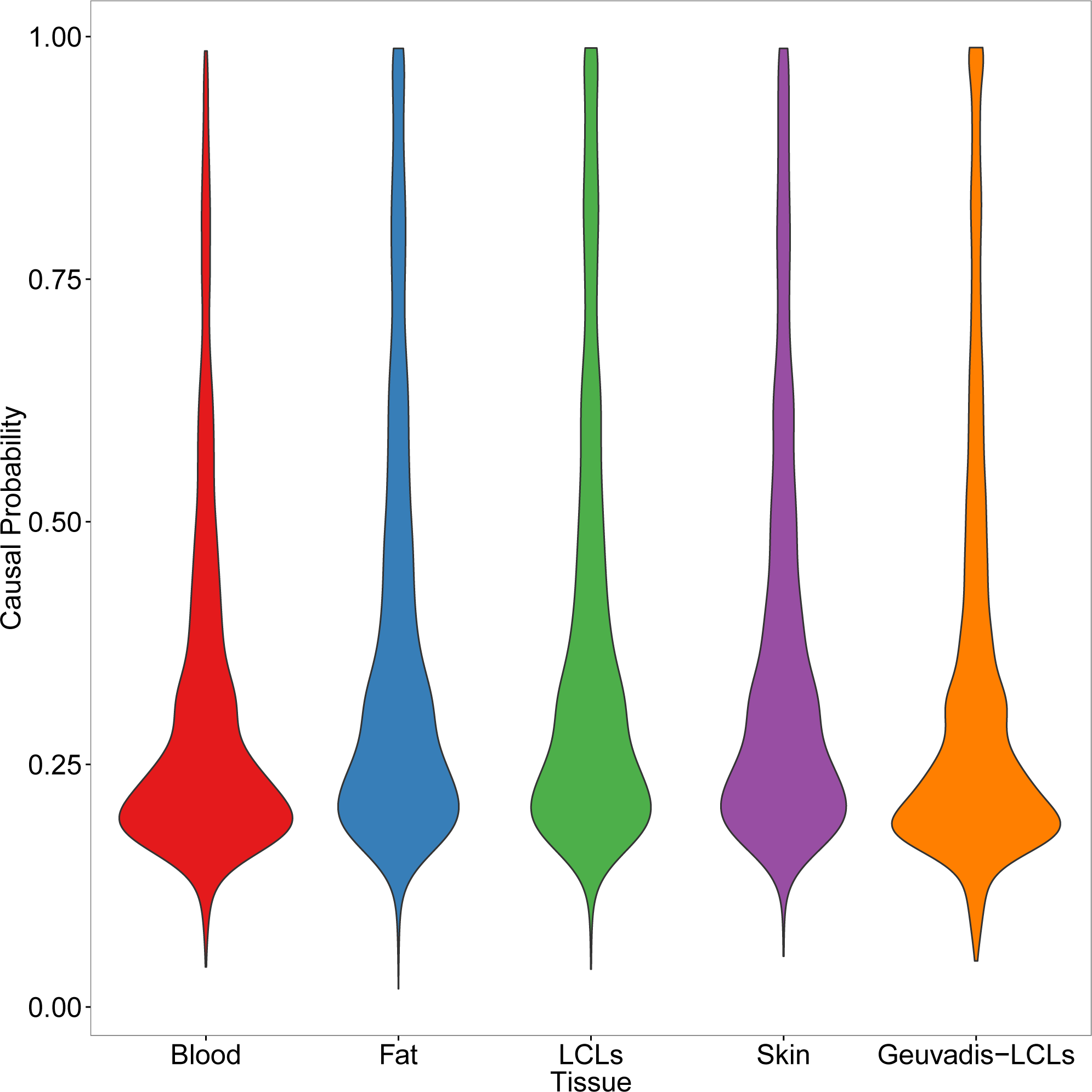
Distribution of the CaVEMaN estimated causal probabilities for all lead eQTLs, broken down by tissue.

To understand more about the relationship between causal regulatory variation and active genomic regions found by ChIP-seq in single individuals, we integrated our causal probabilities with DHSs from the Roadmap Epigenomics consortium. Figure 4 shows a simple linear relationship between the causal probability of the LEV and the probability that the LEV is located in a DHS. We can exploit the linear relationship to estimate the proportion of regulatory variants with causal probability 1 that lie within DHS identified by particular experiments. Figure 5 shows that for all tissues except blood, only a minority of regulatory variants lie within DHS called by specific experiments. Blood eQTL, discovered in a smaller sample size than the other tissues, are more likely to have larger effect sizes and thus affect promoter activity, this is a possible explanation for the observed greater enrichment. It would be interesting to see whether when CaVEMaN is applied to larger eQTL datasets, with the power to discover eQTLs with more subtle effects, the proportion of causal regulatory variants in DHSs will be even lower, implying a limited utility of these regulatory annotations for interpretation of enhancer and weaker regulatory variants.

**Figure 4:**
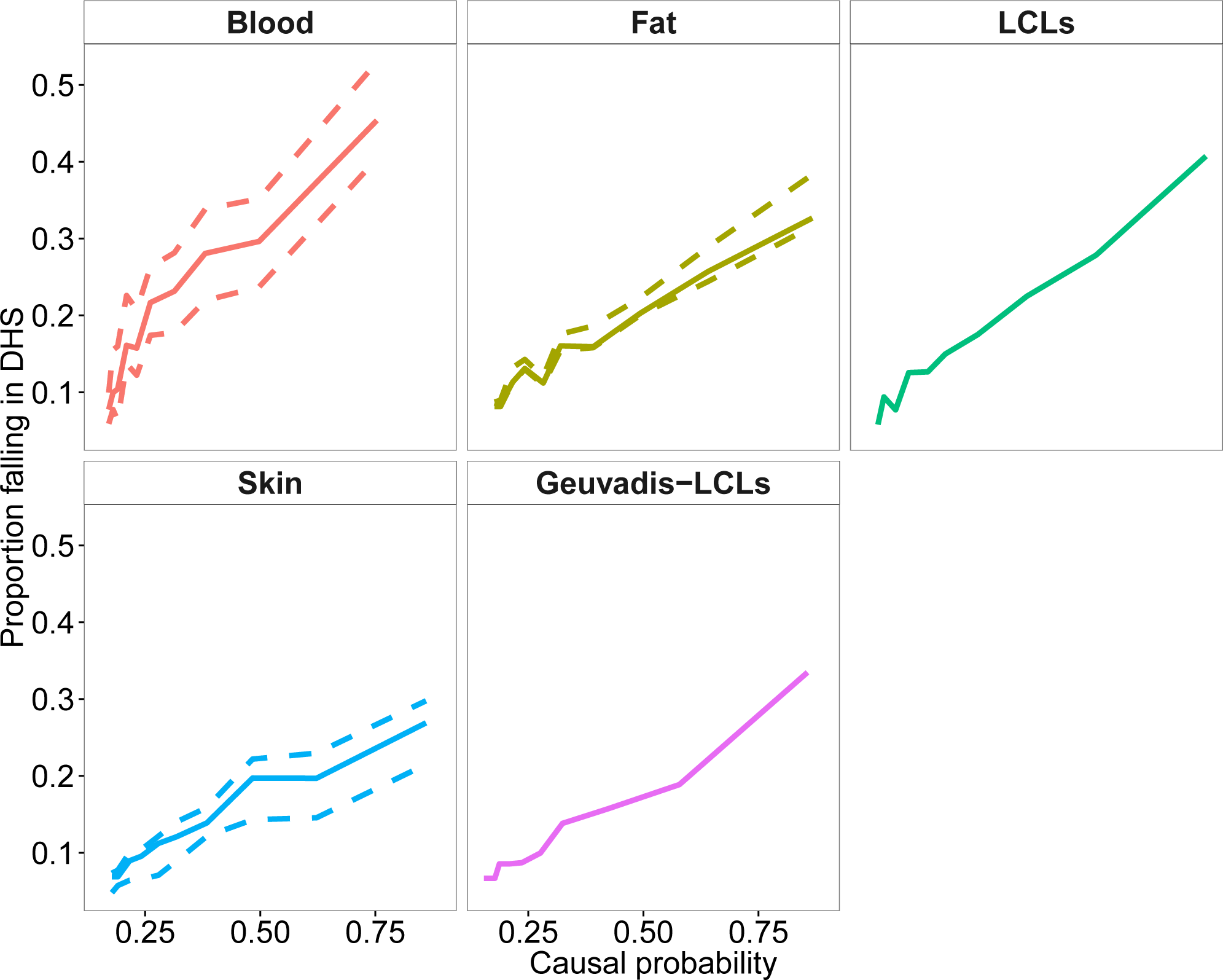
Probability of falling into a DHS is proportion to the CaVEMaN estimated causal probability. The complete line represents the median result across experiments, where there are more than one experiment for a given tissue, the dotted lines give the maximum and minimum across tissues. A full list of experiments can be found in Supplementary Table S2.

**Figure 5:**
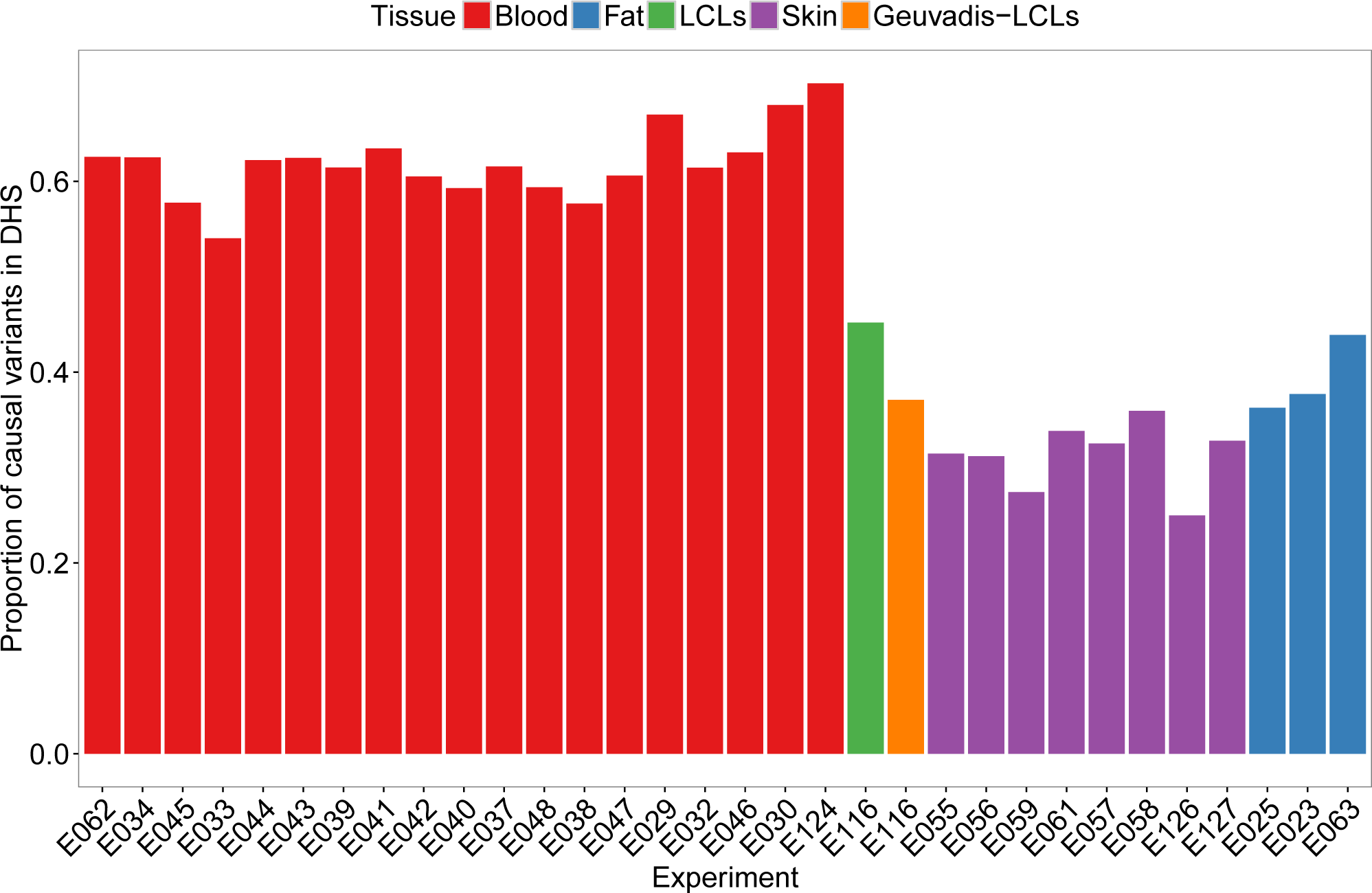
Estimated proportion of functional variants falling into regions identified by single ChIP-seq experiments.

It is widely known that associations with whole organism traits, as discovered by GWAS, are enriched in eQTL (Manolio *et al.*, 2009); by defining a set of eQTL where the causal variant is known we can pinpoint variants which could show greater enrichment (a shared GWAS-eQTL signal would not be diluted by linkage). In addition, by providing both a mediating gene and a variant causative for the expression signal, it is possible these results can provide a more mechanistic understanding of the GWAS signal. By using publicly available GWAS summary statistics from 16 studies (see Supplementary materials), we extracted P values for association for all of the LEVs and saw greater enrichment of small P values for HCCVs compared to all other eQTLs (*π*_1_ = 16.2 compared with *π*_1_ = 14.0, estimated using qvalue (Storey *et al.*, 2015)). Greater enrichment was also observed when considering the proportion of shared signals between GWAS associations with P < 5 × 10^−8^ listed in the NHGRI catalogue and eQTL falling in the same recombination hotspot (16.0% of proximal HCCVs and GWAS associations were shared, compared to 2.49% for all other eQTLs, estimated using the Regulatory Concordance method, RTC, (Nica *et al.*, 2010; Ongen *et al.*, 2016a)). Considering all HCCVs with a Bonferroni significant GWAS association (P < 3 × 10^−6^), we found associations between 53 eQTL and 65 GWAS traits (Figure 6, Supplementary File 2).

**Figure 6:**
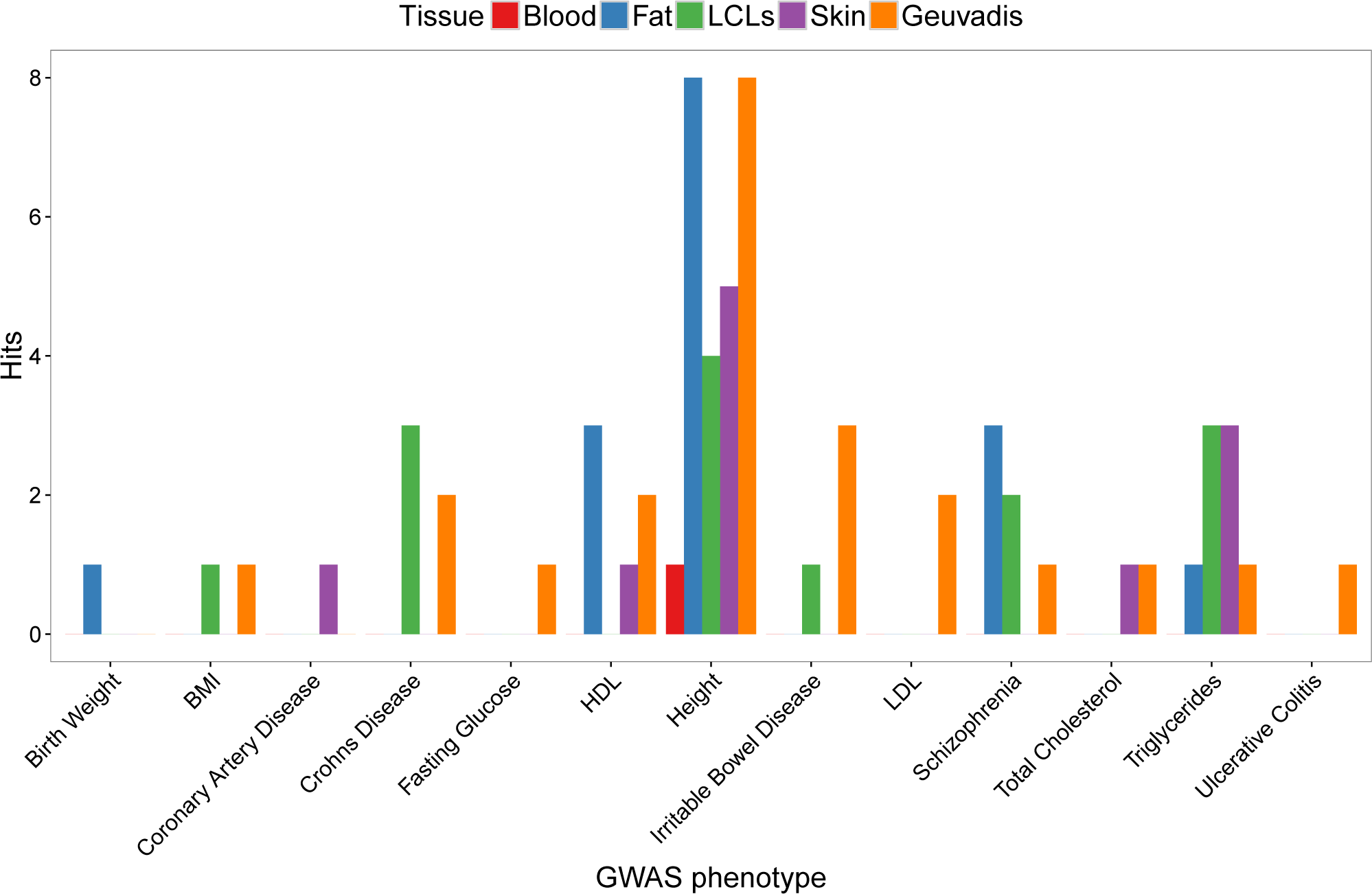
Numbers of significant associations between HCCVs and GWAS traits, divided by tissue type.

Given these examples of variants with highly confident causal effects on expression and statistical associations with GWAS traits, functional evidence connecting the expression of the gene with the trait would also implicate a causal link between variant and trait. For example, a HCCV (rs10274367) associated with *GPER* in is also associated with levels of high-density lipoprotein (HDL) cholesterol. Female knock-out mice for the gene also show a decrease in HDL levels (Sharma *et al.*, 2013). We also found rs1805081 to be both a HCCV for *NPC1*, as well as the lead associated variant with BMI in a large GWAS study (Meyre *et al.*, 2009). Heterozygous mouse models (Npc1+/−), where the gene is expressed at half normal levels, observe large weight gain on high fat diets but not on low fat diets (Jelinek *et al.*, 2010, 2011), and it has also been observed that higher levels of *NPC1* in human adipose tissue normalise after bariatric surgery and behavioural modification (Bambace *et al.*, 2013). In this example, the expression of *NPC1* is modified by rs1805081 and hypothesised to be a response to changes in BMI. Expression changes in *NPC1* then seem to be part of a compensatory mechanism to modify the weight gain due to dietiary excesses and the result of diet-by-genotype interactions. Finally, rs4702 is a HCCV affecting expression of the *FURIN* gene in our analysis and was the lead variant in the GWAS study of schizophrenia (Schizophrenia Working Group of the Psychiatric Genomics, 2014). Following up this association, altering the expression of *FURIN* was seen to produce neuro-anatomical deficits in zebrafish and abnormal neural migration in human induced pluripotent stem cells (Fromer *et al.*, 2016).

In summary, we have produced a method to estimate the probability that the lead eQTL variant is the causal variant. We have used this method to estimate the effectiveness of ChIP-seq experiments from a single individual in predicting regions which harbour regulatory variation, and also to suggest variants which may be causal for GWAS associations. This method could also be applied to GWAS studies, to learn candidate causal variants for whole organism traits. It is clear that pinpointing the causal variant in such studies will not only facilitate the integration of these association signals with mechanistic regulatory interactions and likely upstream regulators, but will also allow the development of interpretation methods from genome sequence alone once a large number of representative causal variants have been discovered.

## Acknowledgments

We would like to thank Nikolaos Lykoskoufis for his help with the enrichment analysis. This work has been supported by grants from the NIH-NIMH (GTEx), European Commission (Direct project), Louis Jeanet Foundation, Swiss National Science Foundation and SystemsX. The TwinsUK study was funded by the Wellcome Trust; European Communitys Seventh Framework Programme (FP7/2007-2013). The study also receives support from the National Institute for Health Research (NIHR)-funded BioResource, Clinical Research Facility and Biomedical Research Centre based at Guy’s and St Thomas’ NHS Foundation Trust in partnership with King’s College London. SNP genotyping was performed by The Wellcome Trust Sanger Institute and National Eye Institute via NIH/CIDR. This study makes use of the data generated by the UK10K Consortium. Funding for UK10K was provided by the Wellcome Trust under award WT091310. A full list of the investigators who contributed to the generation of the data is available at www.UK10K.org. Computation was performed at the Vital-IT (http://www.vital-it.ch) Center for high-performance computing of the SIB Swiss Institute of Bioinformatics.

## Supplementary materials

### TwinsUK data

#### Expression

RPKM expression quantifications used in this paper have been previously analysed (Brown *et al.*, 2014; Buil *et al.*, 2015). In short, eight hundred and fifty-six female twins were recruited from the TwinsUK Adult twin registry and punch biopsies (8 mm) were taken from a photo-protected area adjacent and inferior to the umbilicus. Subcutaneous adipose tissue was separated from skin tissue, and both samples were weighed and immediately stored in liquid nitrogen. Peripheral blood samples were also collected, and the European Collection of Cell Cultures agency generated LCLs by transforming the B-lymphocyte component using the Epstein-Barr virus. The Illumina TruSeq sample preparation kit (Illumina, San Diego, CA) was used to prepare samples according to manufacturer’s instructions, which were then sequenced on a HiSeq2000 machine. The 49-bp sequenced paired-end reads were mapped to the GRCh37 reference genome (Lander *et al.*, 2001) with bwa v0.5.9 (Li and Durbin, 2009). Genes were quantified using the GENCODE v10 annotation (Harrow *et al.*, 2012), and genes defined as protein coding or long non-coding RNA (linc RNA) with less than 10% zero read count were kept. RPKM values were scaled and centred to have mean 0, variance 1 and the first 25 principal components were removed from the whole blood expression and 50 from the other tissues (choice of number of PCs was made a priori based on sample size). Family structure was removed by taking the residuals of an lme4 model (Bates *et al.*, 2014) in which family and zygosity were modelled using random effects. Finally, to remove outlier effects, expression quantifications for each gene were mapped onto a normal distribution with mean 0 and variance 1.

#### Genotyping and genome sequencing

##### Genotypes called from arrays

A combination of the HumanHap300, HumanHap610Q, 1M-Duo and 1.2MDuo Illumina arrays were used to genotype samples. This data was then pre-phased using IMPUTE2 (Howie *et al.*, 2012) and then imputed using the 1000 Genomes Project Phase 1 reference panel (data freeze 10 November 2010, (Abecasis *et al.*, 2012)). For analysis the genotypes were filtered, leaving SNPs with minor allele frequency > 0.01 and IMPUTE info value > 0.8. This data has previously been analysed (Brown *et al.*, 2014; Buil *et al.*, 2015).

##### Genotypes called from sequencing

The vcf files, produced by the UK10K consortium (UK10K Consortium *et al.*, 2015), were downloaded from the European Genome-phenome Archive. When one monozygotic twin in the sample had been sequenced, the same sequence data was used for the genetically identical sibling. Of the 856 individuals with expression, 552 has available sequence data. For multiallelic variants, dosage was calculated as 2 number of copies of the most common allele. Variants were filtered if the major allele had a frequency > 0.99.

##### Ethics statement

The St. Thomas’ Research Ethics Committee (REC) approved on 20 September 2007 the protocol for the dissemination of data, including DNA, with REC reference number RE04/015. On 12 March 2008, the St Thomas’ REC confirmed that this approval extended to expression data. Volunteers gave informed consent and signed an approved consent form before the biopsy procedure. Volunteers were supplied with an appropriate detailed information sheet regarding the research project and biopsy procedure by post before attending for the biopsy. Consent to link the RNA-seq data with the whole genome sequence data was approved by the TwinsUK Resource Executive Committee (TREC) on 22nd April 2015.

#### Geuvadis data

BAM files for the RNA-seq were downloaded from EBI ArrayExpress, accession code E-GEUV-3. These files were mapped to the GRCh37 reference genome (Lander *et al.*, 2001) using GEM version 1.7.1 (Marco-Sola *et al.*, 2012), and protein coding and linc RNAs were quantified using the GENCODE v19 annotation (Harrow *et al.*, 2012). Population group was regressed out of RPKM values as fixed effects in a linear model, values were then centred and scaled to mean 0, variance 1, and 50 principal components were removed. Genotype vcf files from phase 3 of the 1000 Genomes project (1000 Genomes Project Consortium et al. 2015) were downloaded from the 1000 Genomes website. In non-pseudo autosomal regions of the X chromosome, male dosage was calculated as twice the number of copies of the alternate allele (hence treating it as homozygous with two copies). A minor allele frequency cut off of 0.01 was applied.

#### eQTL mapping

eQTLs were mapped using fastQTL (Ongen *et al.*, 2016b). To discover multiple independent eQTLs a stepwise regression procedure was applied. Firstly, for each tissue, fastQTL was run with 10,000 permutations to discover a set of eGenes (FDR < 0.01). Then, the maximum beta-adjusted P value (correcting for multiple testing across the SNPs) over these genes was taken as the gene-level threshold. The next stage proceeded iteratively for each gene. At each iteration a cis scan of the window was performed, using 10,000 permutations and correcting for all previously discovered SNPs. If the beta adjusted P value for the LEV was not significant at the gene-level threshold, the forward stage was complete and the procedure moved on to the backward step. If this P value was significant, the LEV was added to the list of discovered eQTLs as an independent signal and the forward step proceeded to the next iteration.

Once the forward stage was complete for a given gene, a list of associated SNPs was produced which we refer to as forward signals. The backwards stage consisted of testing each forward signal separately, controlling for all other discovered signals. To do this, for each forward signal we ran a cis scan over all variants in the window using fastQTL, fitting all other discovered signals as covariates. If no SNP is significant at the gene-level threshold the signal being tested is dropped, otherwise the LEV from the scan was chosen as the variant that represented the signal best in the full model.

#### Enrichment analysis

Bed files listing DNase hypersensitivity sites, produced by the Roadmap Epigenomics consortium (Roadmap Epigenomics Consortium et al. 2015), were downloaded from the NCBI ftp site). Experiments were linked to tissues from which RNA-seq was available using Table S2. Over each ChIP-seq RNA-seq combination, the odds ratio for enrichment was calculated from the number of LEVs called using sequence and the number of LEVs called using array-based genotypes falling within regions called in the experiment and the total numbers of eQTLs. A Fishers Exact test was performed to test the hypothesis that equal proportions of sequence and genotype LEVs were falling in these regions.

#### Simulations

For all discovered, independent eQTLs, the LEV for association was identified and its minor allele frequency and distance to the transcription start site calculated. In addition, beta and sigma coefficients from a regression of expression on the LEV were also estimated. Then a matched SNP was chosen, with a distance to transcription start site of a protein coding or linc RNA gene within 1 kb of the original, and minor allele frequency within 0.025. Then, simulated expression was produced by multiplying SNP genotype by beta and adding a random normally distributed term with a standard error of sigma. Five simulated datasets were produced for each TwinsUK tissue, eQTL mapping was applied to each looking only for primary eQTLs, and the rank of the nominal P value for association was collected.

#### CaVEMaN

Firstly, we used the simulations to estimate the probability the causal variant would be the ith ranked SNP in an eQTL mapping by calculating the proportion of times this occurred across all tissues and simulations (this quantity is denoted *p_i_*, Supplementary Figure S1). As CaVEMaN focuses on the top 10 ranked variants from an eQTL analysis, *p_i_*, i from 1 to 10, were normalised to sum to 1.

CaVEMaN is based on the premise that there is exactly one genetic signal in the cis window of the gene. For the cases where multiple eQTLs have been discovered for a given gene, we created new single signal expression phenotypes. For each eQTL this was made by regressing out all other eQTLs discovered for the gene, preserving only one genetic signal.

This new matrix of expression data was sampled with replacement 10,000 times to create 10,000 new datasets of the same size. A cis eQTL mapping was run on each of these 10,000 datasets, and the proportion of times a given SNP was ranked i, I from 1 to 10 was calculated (denoted *F_i_*, this is an estimate of the probability that SNP would be the rank i most associated SNP). The CaVEMaN score was then defined as 
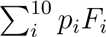
.

Finally, we further exploited the simulations to calibrate the CaVEMaN score of the LEV. CaVEMaN was run on all simulated data. Then, across all simulated datasets (removing blood as this was an outlier resulting in less conservative estimates of causal probabilities) we divided the CaVEMaN scores of the LEVs into twenty quantiles. Within each quantile, we calculated the proportion of times the lead SNP was the causal SNP and then drew a monotonically increasing smooth spline from the origin, through the 20 quantiles, to the point (1, 1) using the gsl interpolate functions with the steffen method (gsl-2.1, Supplementary Figure S2). This function provides our mapping of CaVEMaN score of the lead SNP onto causal probabilities, and we applied this function to the CaVEMaN scores of the LEV to estimate their causal probabilities.

Code for correcting the expression datasets for multiple eQTLs, running the CaVEMaN method and converting the CaVEMaN score to a causal probability can be found here: https://github.com/funpopgen/CaVEMaN.

#### Caviar

For genes with an eQTL in LCLs, we applied Caviar (Hormozdiari *et al.*, 2014) to produce another estimate of causal variant probability for comparison. As Caviar is limited in the number of SNPs it can analyse, we first extracted all variants with P < 0.01, up to the first 50. The Z scores for these variants were produced, with the correlation matrix of these SNPs, and Caviar was run with the default settings.

#### GWAS analysis

We have downloaded the GWAS summary statistics for 16 different GWAS traits: autism (Robinson *et al.*, 2016), birth weight (Horikoshi *et al.*, 2016), body mass index (analysing all ancestries) (Locke *et al.*, 2015), coronary artery disease (Nikpay *et al.*, 2015), Crohns disease (Liu *et al.*, 2015), diabetes (Fuchsberger *et al.*, 2016), fasting glucose (Manning *et al.*, 2012), fasting insulin (Manning *et al.*, 2012), height (Wood *et al.*, 2014), high-density lipoprotein (Global Lipids Genetics Consortium *et al.*, 2013), irritable bowel disease (Liu *et al.*, 2015), low-density lipoprotein (Global Lipids Genetics Consortium *et al.*, 2013), schizophrenia (Schizophrenia Working Group of the Psychiatric Genomics, 2014), total cholesterol (Global Lipids Genetics Consortium *et al.*, 2013), triglycerides (Global Lipids Genetics Consortium *et al.*, 2013), and ulcerative colitis (Liu *et al.*, 2015). For all LEVs, the P value for each trait was extracted (if available) and the qvalue package (Storey *et al.*, 2015) was used to estimate *π*_1_ = 1 − *π*_0_, the proportion of of alternate hypotheses (i.e., association between variant and GWAS trait). Finally, Bonferroni significant GWAS associations for HCCVs were reported, controlling for multiple testing across all phenotypes and variants.

**Table S1:** Estimated costs of collecting whole genome sequence data at GWAS scale relative to expression (WGS is generously priced at $1,000 a genome). Twins UK expression refers to the study published in Buil *et al.* (2015).

| Study | Trait | Sample size | Associations | Estimated Cost* |
| --- | --- | --- | --- | --- |
| GIANT | BMI | 339,224 | 97 | \$339,224,000 |
| PGC | Schizophrenia | 150,064 | 128 | \$150,064,000 |
| MAGIC | Glycemic traits | 133,010 | 53 | \$133,010,000 |
| TwinsUK expression | LCL expression | 814 | 9,555 | \$814,000 |

In addition, we downloaded the NHGRI-EBI Catalog of reported genome-wide significant associations from the EBI website on the 27^th^ September 2016 and removed all with *P* > 5 × 10^−8^ and where the variant was not listed in dbSNP build 148 (Sherry *et al.*, 2001), leaving 11,636 reported associations. RTC, as implemented in QTLtools (Delaneau *et al.*, 2016), was applied with the default settings to look for sharing of these GWAS variants with the discovered eQTLs. As the RTC statistic is uniformly distributed under the null hypothesis of two separate causal loci, independently located within the hotspot, 1 − RTC can be interpreted as a P value for a shared causal variant. The qvalue package (Storey *et al.*, 2015) was then used to estimate *π*_1_, the proportion of GWAS/eQTLs signals in the same recombination interval which were caused by the same underlying variants.

**Figure S1:**
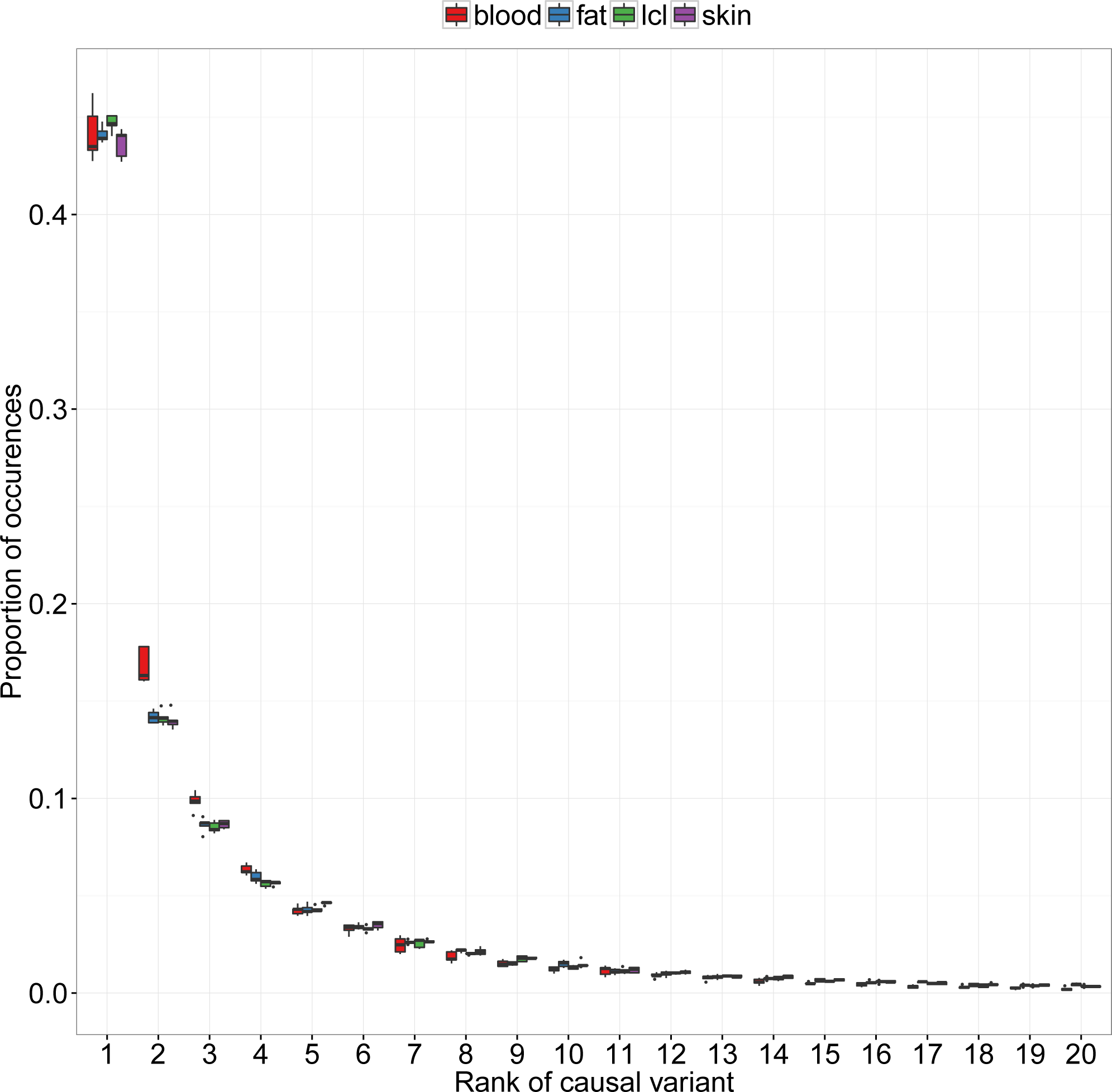
Based on 5 simulations per tissue, the x axis shows the rank of the causal variant, and the y axis the proportion of times this outcome occurred. We notice that, as the whole blood experiment was smaller than the other experiments, sample size does not seem to affect the distribution.

**Figure S2:**
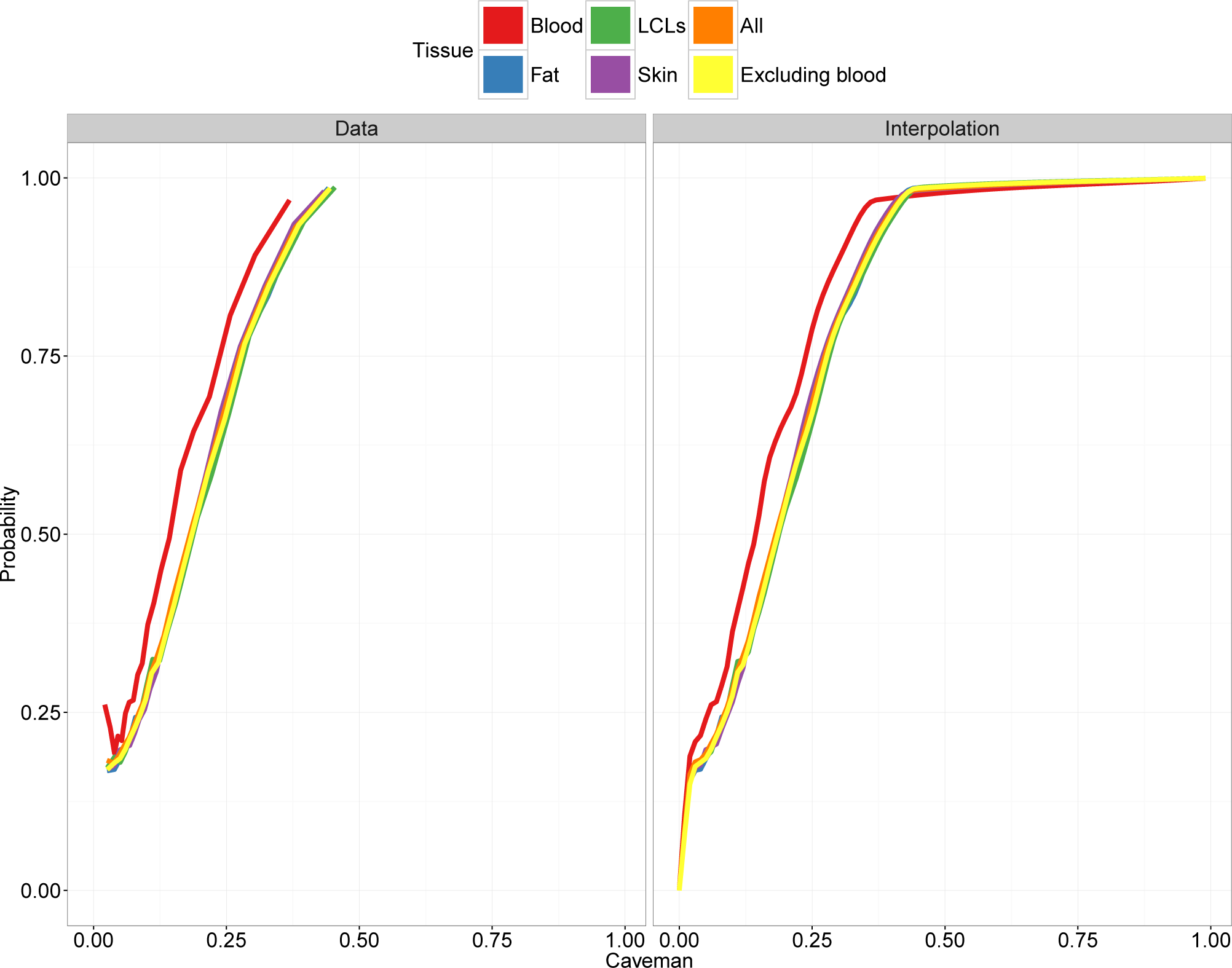
The CaVEMaN score is calibrated using the simulations to estimate the probability that the LEV is causal. The estimated calibration functions are consistent across tissues, with the exception of blood which is less conservative than the other tissues.

**Figure S3:**
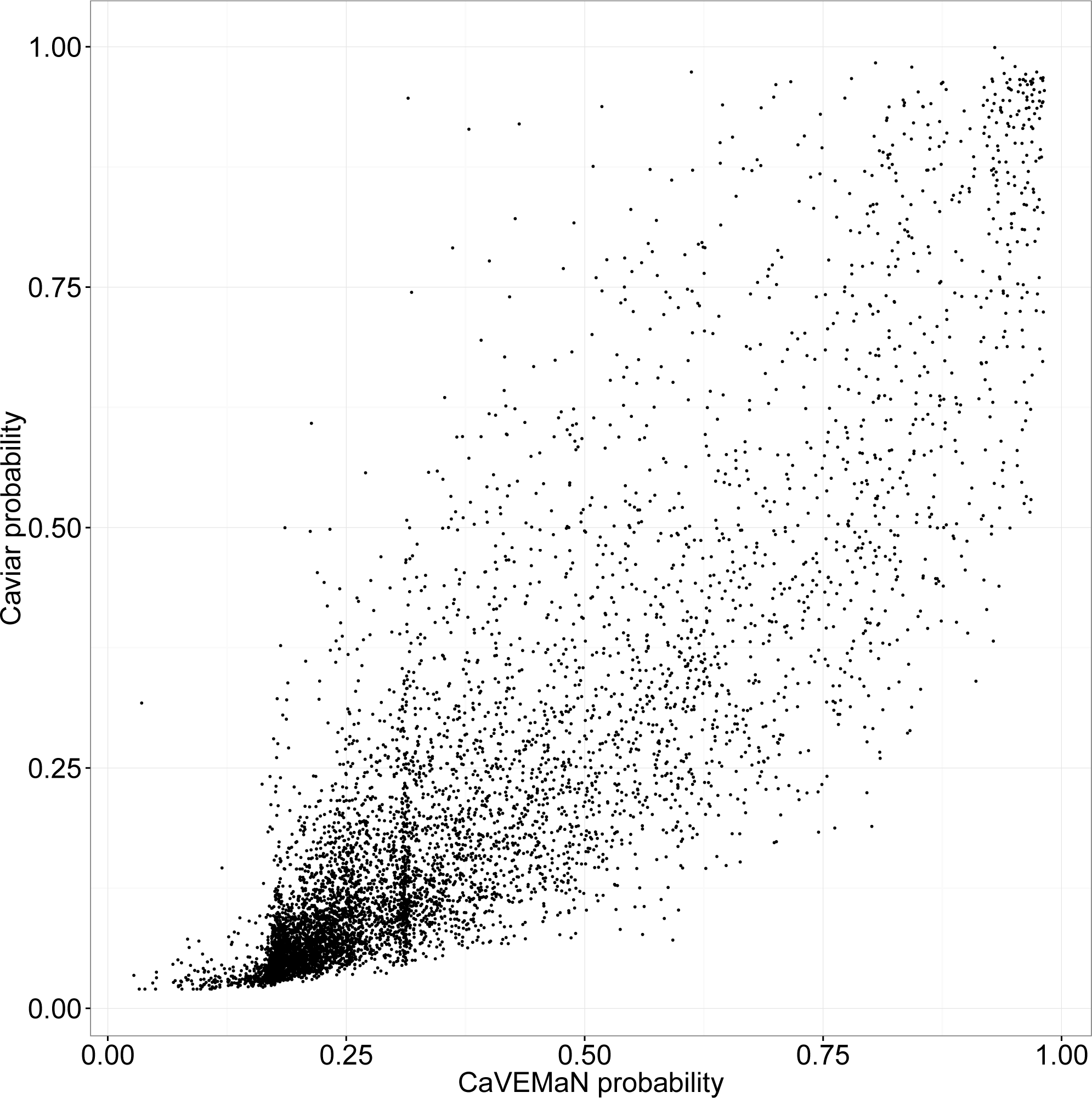
CaVEMaN scores compared to Caviar probabilities for genes with only one eQTL.

**Table S2:** Relevant Roadmap Epigenomics consortium DNAse Hypersensitivity site experiments with code for each analysed RNA-seq experiments. Experiment E116 was used to analyse both TwinsUK and Geuvadis LCLs, all other experiments were specific to one tissue.

| Roadmap Epigenomics experiment | RNA-seq tissue | Roadmap Epigenomics code |
| --- | --- | --- |
| Primary mononuclear cells from peripheral blood | Whole blood | E062 |
| Primary T cells from peripheral blood | Whole blood | E034 |
| Primary T cells effector/memory enriched from peripheral blood | Whole blood | E045 |
| Primary T cells from cord blood | Whole blood | E033 |
| Primary T regulatory cells from peripheral blood | Whole blood | E044 |
| Primary T helper cells from peripheral blood | Whole blood | E043 |
| Primary T helper naive cells from peripheral blood | Whole blood | E039 |
| Primary T helper cells PMA-I stimulated | Whole blood | E041 |
| Primary T helper 17 cells PMA-I stimulated | Whole blood | E042 |
| Primary T helper memory cells from peripheral blood 1 | Whole blood | E040 |
| Primary T helper memory cells from peripheral blood 2 | Whole blood | E037 |
| Primary T CD8+ memory cells from peripheral blood | Whole blood | E048 |
| Primary T helper naive cells from peripheral blood | Whole blood | E038 |
| Primary T CD8+ naive cells from peripheral blood | Whole blood | E047 |
| Primary monocytes from peripheral blood | Whole blood | E029 |
| Primary B cells from peripheral blood | Whole blood | E032 |
| Primary Natural Killer cells from peripheral blood | Whole blood | E046 |
| Primary neutrophils from peripheral blood | Whole blood | E030 |
| Monocytes-CD14+ RO01746 Primary Cells | Whole blood | E124 |
| GM12878 Lymphoblastoid Cells | TwinsUK-LCLs | E116 |
| GM12878 Lymphoblastoid Cells | Geuvadis-LCLs | E116 |
| Foreskin Fibroblast Primary Cells skin01 | Skin | E055 |
| Foreskin Fibroblast Primary Cells skin02 | Skin | E056 |
| Foreskin Melanocyte Primary Cells skin01 | Skin | E059 |
| Foreskin Melanocyte Primary Cells skin03 | Skin | E061 |
| Foreskin Keratinocyte Primary Cells skin02 | Skin | E057 |
| Foreskin Keratinocyte Primary Cells skin03 | Skin | E058 |
| NHDF-Ad Adult Dermal Fibroblast Primary Cells | Skin | E126 |
| NHEK-Epidermal Keratinocyte Primary Cells | Skin | E127 |
| Adipose Derived Mesenchymal Stem Cell Cultured Cells | Subcutaneous adipose | E025 |
| Mesenchymal Stem Cell Derived Adipocyte Cultured Cells | Subcutaneous adipose | E023 |
| Adipose Nuclei | Subcutaneous adipose | E063 |

## References

G. R. Abecasis, A. Auton, L. D. Brooks, M. A. DePristo, R. M. Durbin et al. An integrated map of genetic variation from 1,092 human genomes. Nature, 491(7422):56–65, 2012. doi: 10.1038/nature11632.

C. Bambace, I. Dahlman, P. Arner and A. Kulyté. Npc1 in human white adipose tissue and obesity. BMC Endocrine disorders, 13(1):1, 2013.

D. Bates, M. Mächler, B. Bolker and S. Walker. Fitting linear mixed-effects models using lme4. arXiv preprint arXiv:1406.5823, 2014.

C. Benner, C. C. Spencer, A. S. Havulinna, V. Salomaa, S. Ripatti et al. FINEMAP: efficient variable selection using summary data from genome-wide association studies. Bioinformatics, 32(10):1493–1501, 2016. ISSN 1367-4803. doi:10.1093/bioinformatics/btw018.

A. A. Brown, A. Buil, A. Vinũuela, T. Lappalainen, H.-F. Zheng et al. Genetic interactions affecting human gene expression identified by variance association mapping. eLife, 3:e01,381, 2014. ISSN 2050-084X. doi:10.7554/eLife.01381.

A. Buil, A. A. Brown, T. Lappalainen, A. Vinũuela, M. N. Davies et al. Gene-gene and gene environment interactions detected by transcriptome sequence analysis in twins. Nature Genetics, 47(1):88–91, 2015. doi:10.1038/ng.3162 http://www.nature.com/ng/journal/v47/n1/abs/ng.3162.html\#supplementary-information.

W. Chen, B. R. Larrabee, I. G. Ovsyannikova, R. B. Kennedy, I. H. Haralambieva et al. Fine Mapping Causal Variants with an Approximate Bayesian Method Using Marginal Test Statistics. Genetics, 200(3):719–36, 2015. ISSN 1943-2631. doi:10.1534/genetics.115.176107.

O. Delaneau, H. Ongen, A. A. Brown, A. Fort, N. Panousis et al. A complete tool set for molecular qtl discovery and analysis. bioRxiv, 2016. doi:10.1101/068635.

M. Fromer, P. Roussos, S. K. Sieberts, J. S. Johnson, D. H. Kavanagh et al. Gene expression elucidates functional impact of polygenic risk for schizophrenia. bioRxiv, page 052209, 2016.

C. Fuchsberger, J. Flannick, T. M. Teslovich, A. Mahajan, V. Agarwala et al. The genetic architecture of type 2 diabetes. Nature, 536(7614):41–7, 2016. ISSN 1476-4687. doi:10.1038/ nature18642.

Global Lipids Genetics Consortium, C. J. Willer, E. M. Schmidt, S. Sengupta, G. M. Peloso et al. Discovery and refinement of loci associated with lipid levels. Nature genetics, 45(11):1274–83, 2013. ISSN 1546-1718. doi:10.1038/ng.2797.

J. Harrow, A. Frankish, J. M. Gonzalez, E. Tapanari, M. Diekhans et al. GENCODE: the reference human genome annotation for The ENCODE Project. Genome Res, 22(9):1760–1774, 2012. doi:10.1101/gr.135350.111.

M. Horikoshi, R. N. Beaumont, F. R. Day, N. M. Warrington, M. N. Kooijman et al. Genome-wide associations for birth weight and correlations with adult disease. Nature, doi:10.1038/nature19806, 2016.

F. Hormozdiari, E. Kostem, E. Y. Kang, B. Pasaniuc and E. Eskin. Identifying causal variants at loci with multiple signals of association. Genetics, 198(2):497–508, 2014. ISSN 1943-2631. doi:10.1534/genetics.114.167908.

B. Howie, C. Fuchsberger, M. Stephens, J. Marchini and G. R. Abecasis. Fast and accurate genotype imputation in genome-wide association studies through pre-phasing. Nat Genet, 44(8):955–959, 2012. doi:10.1038/ng.2354.

D. Jelinek, R. A. Heidenreich, R. P. Erickson and W. S. Garver. Decreased npc1 gene dosage in mice is associated with weight gain. Obesity, 18(7):1457–1459, 2010.

D. Jelinek, V. Millward, A. Birdi, T. P. Trouard, R. A. Heidenreich et al. Npc1 haploinsufficiency promotes weight gain and metabolic features associated with insulin resistance. Human molecular genetics, 20(2):312–321, 2011.

E. S. Lander, L. M. Linton, B. Birren, C. Nusbaum, M. C. Zody et al. Initial sequencing and analysis of the human genome. Nature, 409(6822):860–921, 2001. doi:10.1038/35057062.

T. Lappalainen, M. Sammeth, M. R. Friedlander, P. A. t Hoen, J. Monlong et al. Transcriptome and genome sequencing uncovers functional variation in humans. Nature, 501(7468):506–511, 2013. doi:10.1038/nature12531.

C. M. Lebreton and P. M. Visscher. Empirical nonparametric bootstrap strategies in quantitative trait loci mapping: conditioning on the genetic model. Genetics, 148(1):525–35, 1998. ISSN 0016-6731.

H. Li and R. Durbin. Fast and accurate short read alignment with Burrows-Wheeler transform. Bioinformatics, 25(14):1754–1760, 2009. doi:10.1093/bioinformatics/btp324.

J. Z. Liu, S. van Sommeren, H. Huang, S. C. Ng, R. Alberts et al. Association analyses identify 38 susceptibility loci for inflammatory bowel disease and highlight shared genetic risk across populations. Nature genetics, 47(9):979–86, 2015. ISSN 1546-1718. doi:10.1038/ng.3359.

A. E. Locke, B. Kahali, S. I. Berndt, A. E. Justice, T. H. Pers et al. Genetic studies of body mass index yield new insights for obesity biology. Nature, 518(7538):197–206, 2015. doi: 10.1038/nature14177.

A. K. Manning, M.-F. Hivert, R. A. Scott, J. L. Grimsby, N. Bouatia-Naji et al. A genomewide approach accounting for body mass index identifies genetic variants influencing fasting glycemic traits and insulin resistance. Nature genetics, 44(6):659–69, 2012. ISSN 1546-1718. doi:10.1038/ng.2274.

T. A. Manolio, F. S. Collins, N. J. Cox, D. B. Goldstein, L. A. Hindorff et al. Finding the missing heritability of complex diseases. Nature, 461(7265):747–753, 2009. doi:10.1038/nature08494.

J. Marchini and B. Howie. Genotype imputation for genome-wide association studies. Nat Rev Genet, 11(7):499–511, 2010. doi:10.1038/nrg2796.

S. Marco-Sola, M. Sammeth, R. Guigo and P. Ribeca. The GEM mapper: fast, accurate and versatile alignment by filtration. Nat Methods, 9(12):1185–1188, 2012. doi:10.1038/nmeth. 2221.

D. Meyre, J. Delplanque, J.-C. Chèvre, C. Lecoeur, S. Lobbens et al. Genome-wide association study for early-onset and morbid adult obesity identifies three new risk loci in european populations. Nature genetics, 41(2):157–159, 2009.

A. C. Nica, S. B. Montgomery, A. S. Dimas, B. E. Stranger, C. Beazley et al. Candidate causal regulatory effects by integration of expression QTLs with complex trait genetic associations. PLoS Genet, 6(4):e1000,895, 2010. doi:10.1371/journal.pgen.1000895.

M. Nikpay, A. Goel, H.-H. Won, L. M. Hall, C. Willenborg et al. A comprehensive 1,000 Genomes-based genome-wide association meta-analysis of coronary artery disease. Nature genetics, 47(10):1121–30, 2015. ISSN 1546-1718. doi:10.1038/ng.3396.

H. Ongen, A. A. Brown, O. Delaneau, N. Panousis, A. C. Nica et al. Estimating the causal tissues for complex traits and diseases. bioRxiv, page 074682, 2016a.

H. Ongen, A. Buil, A. A. Brown, E. T. Dermitzakis and O. Delaneau. Fast and efficient QTL mapper for thousands of molecular phenotypes. Bioinformatics, 32(10):1479–1485, 2016b. ISSN 1367-4803. doi:10.1093/bioinformatics/btv722.

Roadmap Epigenomics Consortium, A. Kundaje, W. Meuleman, J. Ernst, M. Bilenky et al. Integrative analysis of 111 reference human epigenomes. Nature, 518(7539):317–30, 2015. ISSN 1476-4687. doi:10.1038/nature14248.

E. B. Robinson, B. St Pourcain, V. Anttila, J. A. Kosmicki, B. Bulik-Sullivan et al. Genetic risk for autism spectrum disorders and neuropsychiatric variation in the general population. Nature genetics, 48(5):552–5, 2016. ISSN 1546-1718. doi:10.1038/ng.3529.

C. Schizophrenia Working Group of the Psychiatric Genomics. Biological insights from 108 schizophrenia-associated genetic loci. Nature, 511(7510):421–427, 2014. doi:10.1038/ nature13595.

B. Servin and M. Stephens. Imputation-based analysis of association studies: candidate regions and quantitative traits. PLoS genetics, 3(7):e114, 2007. ISSN 1553-7404. doi:10.1371/journal.pgen.0030114.

G. Sharma, C. Hu, J. L. Brigman, G. Zhu, H. J. Hathaway et al. Gper deficiency in male mice results in insulin resistance, dyslipidemia, and a proinflammatory state. Endocrinology, 154(11):4136–4145, 2013.

S. T. Sherry, M.-H. Ward, M. Kholodov, J. Baker, L. Phan et al. dbsnp: the ncbi database of genetic variation. Nucleic acids research, 29(1):308–311, 2001.

S. L. Spain and J. C. Barrett. Strategies for fine-mapping complex traits. Human molecular genetics, 24(R1):R111–R119, 2015.

J. Storey, A. Bass, A. Dabney and D. Robinson. qvalue: Q-value estimation for false discovery rate control, 2015. R package version 2.2.2.

UK10K Consortium, K. Walter, J. L. Min, J. Huang, L. Crooks et al. The UK10K project identifies rare variants in health and disease. Nature, 526(7571):82–90, 2015. ISSN 1476-4687. doi:10.1038/nature14962.

P. M. Visscher, R. Thompson and C. S. Haley. Confidence intervals in QTL mapping by bootstrapping. Genetics, 143(2):1013–20, 1996. ISSN 0016-6731.

D. Welter, J. MacArthur, J. Morales, T. Burdett, P. Hall et al. The NHGRI GWAS Catalog, a curated resource of SNP-trait associations. Nucleic Acids Res, 42(Database issue):D1001–6, 2014. doi:10.1093/nar/gkt1229.

A. R. Wood, T. Esko, J. Yang, S. Vedantam, T. H. Pers et al. Defining the role of common variation in the genomic and biological architecture of adult human height. Nat Genet, 46(11):1173–1186, 2014. doi:10.1038/ng.3097.

